# Signals in flux: Investigating the seasonal turnover of the diel vertical migration-inducing kairomone 5α-cyprinol sulfate

**DOI:** 10.1101/2025.07.07.663427

**Authors:** Johanna Ahlers, Eric von Elert

**Affiliations:** Aquatic Chemical Ecology, Department of Biology, University of Cologne, Zülpicher Str.47B, 50674 Köln, Germany

**Keywords:** DVM, *Daphnia*, Kairomones, Microbial degradation, Chemical ecology

## Abstract

5α-cyprinol sulfate (CPS) is a chemical signal released by cyprinid fish that plays a crucial role in predator-prey interactions by inducing diel vertical migration (DVM) in *Daphnia*, a key process driving freshwater ecosystem dynamics. While CPS-mediated DVM has long been considered a stable and predictable response to predator abundance, this study challenges previous assumptions by demonstrating that CPS undergoes rapid turnover in aquatic systems. We show that CPS exudation is tightly linked to fish feeding intensity, independent of fish size, while microbial degradation emerges as the dominant removal process, with degradation rates reaching up to 85.7% per day in lake water. Seasonal field data from a mesotrophic lake reveal that CPS concentrations range from 40 to 900 ng/L over a season, peaking in late summer, putatively indicating increased fish feeding and shifts in prey availability. Using an integrated approach combining laboratory feeding trials, field sampling, and predictive modeling, we present a temperature-dependent framework that captures the interplay between CPS exudation and degradation. Our findings highlight a fundamental reconsideration of CPS as an environmental cue: rather than reflecting predator abundance, CPS concentrations primarily track predator feeding activity, dynamically modulating the strength of DVM and its cascading effects on freshwater food webs. This study provides the first in-situ quantification of CPS turnover, offering new insights into the temporal variability of chemical signaling and its ecosystem-wide consequences in fluctuating aquatic environments.

## Introduction

Chemical signaling is a fundamental mechanism driving ecological interactions in aquatic systems, shaping community structures and sustaining ecosystem functions (von Elert, 2012). Among the diverse organisms responding to such signals, *Daphnia*, a genus of freshwater planktonic crustaceans, plays a crucial role in aquatic food webs as a primary grazer and a key prey species for fish (Miner et al. 2012). To cope with predation, *Daphnia* exhibit a variety of inducible defenses, including behavioral shifts (e.g., diel vertical migration (DVM)), morphological changes, and life-history modifications. These adaptive responses enhance predator avoidance and improve survival chances. Among the signals triggering these defenses, 5α-cyprinol sulfate (CPS) has recently emerged as a key kairomone released by cyprinid fish, inducing defensive traits in *Daphnia* (Hahn et al. 2019; Hahn and von Elert 2022; Ahlers and Hahn et al. 2024) Thus, CPS plays a pivotal role in predator-prey interactions and broader ecological dynamics in aquatic environments.

From a chemical and environmental perspective, the bile salt CPS possesses properties that influence its behavior in natural systems. As a low-molecular-weight (<500 Da), non-volatile, anionic compound of medium polarity, CPS exhibits remarkable chemical stability across extreme pH levels (0.8–14.0) and temperatures (−20°C to +120°C) (Loose et al. 1993; von Elert and Loose 1996). This high stability is believed to have evolutionary significance, making bile compounds in general and CPS in particular well-suited to function as infochemicals (Buchinger et al. 2014), as it is reasonable to assume that this resilience enables bile compounds to persist and remain effective under diverse and challenging environmental conditions. Additionally, CPS’s amphiphilic nature theoretically allows it to form micelles, which could enhance its dispersal and chemical stability in aquatic environments (von Elert and Pohnert 2000; Goto et al. 2003).

However, potential degradation of CPS in natural water bodies remains largely unexplored, but insights from other low molecular weight organic compounds suggest that microbial and photochemical processes may play an important role (Sterr and Sommaruga 2008; Ghuneim et al. 2021). Based on studies of other kairomones, CPS degradation by microbes can been suggested. Loose et al. (1993) demonstrated removal of DVM-inducing activity upon incubation with bacteria and hypothesized that similar processes may occur in natural water bodies, limiting the persistence of kairomones in aquatic systems. This idea was supported by Beklioglu et al. (2006), who demonstrated the rapid disappearance of DVM of *Daphnia* in response to cyprinid fish water in laboratory conditions. While these studies highlight plausible microbial degradation, no direct evidence currently exists to confirm whether CPS undergoes microbial breakdown under environmentally relevant conditions. Importantly, for a kairomone to serve as a reliable indicator of current predation risk, it must degrade in a timely manner once the threat subsides. If such signals were too persistent, prey organisms might continue investing in costly defensive traits even when they are no longer necessary, leading to reduced fitness. Therefore, the degradation of CPS is not only ecologically relevant but also essential for the adaptive value of predator-induced responses.

Temperature is a critical factor influencing many biological and chemical processes in aquatic ecosystems, including fish exudation and microbial degradation. Studies show that temperature-dependent excretion rates in fish are linked to metabolic activity and food intake (Persson 1982; Hölker 2003; Hölker 2006). However, whether CPS exudation of fish follows similar temperature-related patterns remains untested. Similarly, microbial activity accelerates the degradation of organic compounds, including bioactive substances (Simon and Wünsch 1998; Vrede 2005), but the impact of temperature on CPS degradation by microbial communities remains unknown. The dynamic interplay between CPS exudation and degradation likely determines its persistence and signaling efficacy in natural systems. Warmer temperatures may enhance both exudation and degradation, potentially stabilizing CPS concentrations if the processes counterbalance. Alternatively, if degradation outpaces exudation under conditions of high microbial activity and low fish density or reduced feeding intensity, CPS levels might decline. Conversely, high exudation rates during periods of intensive feeding and elevated metabolic activity of fish (e.g., during zooplankton blooms in summer) could sustain CPS levels despite increased degradation. This highlights the critical role of temperature in modulating CPS turnover, yet the extent of these effects remains poorly understood.

To address these knowledge gaps, this study investigates whether CPS turnover is temperature-dependent by examining both exudation and degradation processes through an integrated approach of laboratory experiments, field observations, and mathematical modeling. Specifically, we aim to (i) determine how fish size, food intake, and temperature influence CPS release; (ii) investigate the role of temperature on microbial CPS breakdown and (iii) examine temporal fluctuations of in-situ CPS concentrations and their relationship with environmental factors. By integrating experimental data and modeling, we aim to elucidate the biotic (fish exudation, microbial degradation) and abiotic (temperature) drivers shaping CPS dynamics. This study offers new insights into the ecological significance of CPS in predator-prey interactions and ecosystem stability under changing environmental conditions.

## Materials and Methods

### In situ assessment of CPS degradation in lake water

To investigate the in-situ degradation rates of 5-α-cyprinol sulfate (CPS), we collected water samples in June 2024 from the epilimnion (1–2 m depth) of two lakes: Reeser Meer Nord Erweiterung (RMNE) (51°45’3” N 6°27’59” E) and Fühlinger See (FS) (51°1’23” N 6°55’27” E). RMNE covers an area of 0.618 km², with a maximum depth of 10.5 m and was originally formed by gravel excavation and has undergone extensive restoration efforts. Populations of the cyprinid species *Leucaspius delineatus* were first recorded in RMNE in 2019 (Jost Borcherding, pers. comm.). The phosphorus concentration in the lake water, measured as total phosphorus (TP), indicated that RMNE is mesotrophic (mean TP value of 13 µg P/L across the water column). This measurement was conducted using a segmented flow analyzer (SKALAR, Breda, The Netherlands) and a wavelength of 880 nm (DIN EN ISO 15681 Part 2-D46). FS spans an area of 1 km² and reaches a maximum depth of 14 m (www.fuehlinger-see-koeln.de). Parts of the lake are designated for recreational use. The cyprinid species present in FS include Bream (*Abramis brama*), Tench (*Tinca tinca*), Common Roach (*Rutilus rutilus*), Rudd (*Scardinius erythrophthalmus*), and Common Carp (*Cyprinus carpio*) (www.asg-ford.com). The phosphorus analysis for FS also indicated a mesotrophic status (mean TP value of 14 µg P/L across the water column).

Using a Ruttner water sampler, water was collected from FS on June 4, 2024, and from RMNE on June 12, 2024. The samples were then filtered through a 250 µm plankton net to remove large organisms. Filtered water was stored in 1 L glass bottles, with four liters subsequently being autoclaved to eliminate microbial activity (Control) and four untreated (NA treatment). The bottles were incubated at 18°C, matching in-situ temperatures at the time of the sampling. After 20 hours, aliquots of 200 mL were removed under sterile conditions from each of the bottles. Then, each aliquot was filtered through a glass microfiber filter (Whatman Grade GF/A: 1.6 μm) to remove particles. To each filtered sample, we added an internal standard (100 ng of cholesteryl sulfate) to account for variability during sample processing. Additionally, 2 mL of 100% methanol was added to each sample to prepare them for solid-phase extraction (SPE). The samples were then subjected to C18-SPE cartridges (as detailed below). To determine the degradation rate of CPS, this procedure was repeated daily for four consecutive days. Daily CPS concentrations from each of the replicate bottles resulted in scatter plots of CPS concentration over time for Control and NA treatment for both lakes. An exponential decay curve was then fitted to these data points to estimate the degradation rate constant (k) (as detailed below).

### Assessment of CPS exudation rates in fish

To assess the exudation rates of CPS, individual roach fish (*Rutilus rutilus*) were kept at 18 °C in separate, darkened aquaria filled with 20 L tap water. Fish were categorized as small and large, and showed a significant difference in body mass (Welch’s *t*-test, p < 0.001). Small fish weighed 18.2 ± 6.7 g (range = 12–29 g, n = 13), whereas large fish weighed 89.9 ± 13.5 g (range = 77–110 g, n = 14). Prior to the experiment, fish were acclimated without food in their respective aquaria for three days to standardize conditions and ensure empty digestive tracts. At the start of the experiment, individual fish were fed varying amounts of food based on their body weight. Fish were fed frozen chironomid larvae (www.nature2aqua.de, Article no. 624010). To simulate a natural feeding regime and avoid overfeeding during the short feeding period, fish received only 1/6 of their daily ration, corresponding to 4 hours of the feeding cycle. This ration, when scaled to a 24-hour period, would constitute 0.5–5% of the fish’s body weight. All of the provided food was fully consumed by the fish within the 4-hour feeding period. Following the feeding period, fish were transferred into 20 L of fresh tap water for a 20-hour incubation. Then, two 1 L water samples were taken from each aquarium to serve as technical replicates. Water samples were filtered through glass microfiber filter (Whatman Grade GF/A: 1.6 μm) to remove particles. To each 1 L sample, 500 ng of cholesteryl sulfate was added as an internal standard. Subsequently, 10 mL of 100% methanol was added to each sample, which then was subjected to C18-SPE (as detailed below). To determine total CPS exudation per fish, CPS exudation was totaled over four consecutive days following feeding.

### Seasonal CPS concentrations in lake Reeser Meer Nord Erweiterung (RMNE)

To assess seasonal changes of in-situ CPS concentrations in RMNE, surface water samples were collected using a bucket. Samples were collected monthly during the first or second week of each month from March to September 2023. The water samples were immediately transported to the laboratory and then filtered through glass microfiber filters (Whatman Grade GF/A: 1.6 μm). Four 1 L subsamples (technical replicates) were taken from the filtered water, and 500 ng of cholesteryl sulfate was added as an internal standard to each replicate for normalization during quantification. The samples were stored overnight at 7–10°C to maintain stability and minimize degradation processes, because immediate extraction was logistically infeasible. The other day, 10 mL of 100% methanol was added to each sample to prepare them for C18-SPE (detailed below).

### Solid phase extraction

The C18-SPE cartridges (Agilent Bond Elute C18, 500 mg, 3 ml) were activated with 10 ml of 100% methanol (MeOH) and subsequently conditioned with 1% MeOH dilution. Each sample was drawn into the cartridge at a controlled flow rate of approximately 50 µL/s using silicone tubing, ensuring a stable flow and minimizing disturbances during the extraction process. The samples were then washed with 10 mL 1% MeOH and finally eluted with 10 mL of 100% MeOH, collected in a glass tube. Each sample was applied to a new cartridge to avoid cross-contamination. Technical replicates were applied to the same cartridges. The collected samples were dried overnight in a vacuum centrifuge at 30 °C and 1500 rpm. Once dried, samples were re-dissolved in 200 μL of 100% MeOH and transferred into glass vials for further analysis.

### Mass spectrometry and quantification of CPS

Mass analysis was conducted by applying 5 µl of each sample to a Vanquish™ Horizon UHPLC system coupled with a Q-Exactive HF mass spectrometer (Thermo Fisher Scientific, Dreieich, Germany). Metabolite separation was achieved on an EC 125/2 Nucleosil 100-3 C18 column (Macherey-Nagel, Düren, Germany) maintained at 30 °C and a constant flow rate of 300 µL/min. The mobile phase comprised solvent A (13 mM ammonium acetate and 10 mM formic acid in water) and solvent B (0.01 % ammonium acetate and 10 mM formic acid in a 9:6 [v/v] mixture of acetonitrile and methanol). The following gradient programs were applied: Gradient A (20% B, 0.0–28 min gradient to 100% B; 28–32 min at 100% B; 32–33 min gradient to 20% B; 33–35 min at 20% B) and Gradient B (30% B, 0.0–12 min gradient to 100% B; 12–14 min at 100% B; 14–15 min gradient to 30% B; 15–17 min at 30% B). We used two different gradient programs because a longer gradient (Gradient A) was chosen for the seasonal lake water samples from RMNE to potentially detect additional seasonal patterns for other kairomones in future analyses. Negative and positive ionization was performed for each run simultaneously, with a spray voltage of 3.0–3.53 kV and a capillary temperature set to 360 °C. To ensure accurate CPS quantification, a calibration curve for the internal standard cholesteryl sulfate was included in each sample sequence.

### Modeling the temperature-dependent release of CPS

From the perspective of an individual cyprinid fish, the net release of CPS normalized to the food ration is the difference between the rate of exudation from the individual fish and the rate of microbial degradation. Since the exudation from the individual fish proved to scale linearly with food ration (see Results), we here model the net release of CPS normalized to the food ration. To model the net release of CPS into the aquatic environment, we developed a temperature-dependent approach that integrates CPS exudation rates and degradation rates. The model assumes a dynamic balance between the exudation of CPS by fish and its removal by microbial activity (degradation). The temperature-dependent exudation rate (*E*) and degradation rate (*D*) are calculated using Q10 scaling, which accounts for the respective exponential effect of temperature on these biological processes. Specifically, the exudation rate is adjusted from a reference rate (*E_ref_*) measured at a reference temperature (*T_ref_*) for a given food ration using the equation

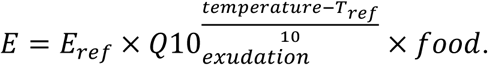

Similarly, the degradation rate is adjusted from a reference rate (D_ref_) using

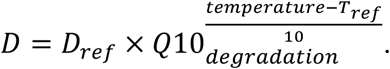

The ratio of the rate of exudation E and the rate of degradation D gives the net release of CPS in ng/fish

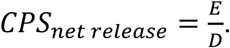

This model enabled us to simulate the net CPS release across a range of temperatures, providing insights into the interplay between fish-mediated exudation and microbial degradation under different temperatures. The parameter values for the reference rates (E_ref_ = 63.68 ng/h x gram of food per fish and D_ref_ ∼ 0.0778 1/h) were derived from experimental data. Q10 factors were calculated based on the standard metabolic rate of roach (*Rutilus rutilus*), yielding a value of 2.4 (Hölker 2006), and bacterial cell multiplication, yielding a value of ∼2 (Simon and Wünsch 1998). This model is an equilibrium-based simplification and assumes that the system is in a dynamic steady state—where inputs and outputs are ongoing but remain balanced. The model focuses on the long-term average CPS concentration in the water under stable conditions of temperature, exudation, and degradation, rather than capturing transient fluctuations or short-term changes. To demonstrate how this model can be applied, we incorporated predicted daily in-situ feeding rations of cyprinids based on Persson (1983). This example illustrates how the model can help predict CPS dynamics in other aquatic ecosystems and serves as a framework to obtain information about the potential ecological impacts of CPS-mediated signaling.

**Exudation Rate (E):** The reference exudation rate (E_ref_) is defined as the amount of CPS exuded per hour per gram of food consumed per fish, expressed in ng/h/fish. This rate represents the amount of CPS exuded into the water under reference conditions (18°C). The temperature dependence of exudation is modeled using a Q_10-_factor, which scales E with temperature. The experimental results determined a reference exudation rate (E_ref_) of 64.5 ng/h x gram of food per fish. This rate was determined using a linear regression model analyzing the relationship between final CPS concentration in the water and food amount consumed. Fish size was omitted from the model, as it was found to have no significant effect on the CPS concentration released by the fish.

**Degradation Rate (D):** The degradation rate (D) is defined as the rate at which a compound degrades. It is determined using the degradation constant k derived from an exponential decay model. In this model, the degradation follows first-order kinetics, meaning the rate of degradation is proportional to the concentration of the compound. The rate constant k is influenced by temperature, with higher temperatures generally accelerating degradation. This temperature dependence is incorporated using a Q_10_**-**factor, which scales k with temperature changes. For the purpose of this model, we focus on a degradation rate of approximately 7,78% per hour, corresponding to a reference degradation rate (D_ref_) of 0.0778 per hour under reference conditions (18°C), based on our experimental data for CPS degradation in RMNE.

### Zooplankton sampling

Zooplankton samples in RMNE were collected once per month from March to September 2023 at two time points per day: midday (10 am–3 pm) and night (9 pm–11 pm). At each time point, five samples were taken from both the epilimnion (∼2 m) and hypolimnion (∼9 m) using a Schindler-Patalas trap (10 L capacity and a 200 μm mesh size). This resulted in a total of 20 samples per month. Samples were preserved in a sucrose-ethanol solution (285 mL distilled water, 714 mL ethanol (C₂H₆O) (96–98%), 40 g sucrose (C₁₂H₂₂O₁₁), and 40 mL glycerol (C₃H₈O₃)) and were analyzed using a HydroptiC ZooScan system (Vogelmann et al. 2022) and Plankton Identifier (PkID) software (Gorsky et al. 2010). Classification was conducted semi-automatically based on a validated learning set, and *Daphnia* abundance was calculated as the sum of all samples across both depths and both time points (Jost Borcherding, pers. comm.).

### Statistical analysis

All statistical analyses were conducted in R (version 4.2.0; R Core Team, 2022) within the RStudio environment (Posit Team, 2023) to explore CPS degradation, exudation dynamics, and model predictions. For degradation experiments, a linear mixed-effects model (lme) was applied to analyze log-transformed CPS concentrations over time. This approach linearized the exponential decay relationship and allowed for the inclusion of replicates as random factors, accounting for variability between experimental replicates. The lme model facilitated the estimation of degradation rate constants (k) and initial CPS concentrations (C_0_) for different treatments, while also testing for statistically significant differences between lakes and treatments. Temperature-dependent degradation rates at 15°C and 25°C were predicted using the Q10 scaling factor, derived from the estimated rate constants at 18°C. For exudation experiments, linear regression models were applied to assess the relationships between CPS exudation rates, food intake, and fish size, highlighting the critical influence of feeding activity on exudation dynamics. Furthermore, predictive modeling incorporated Q10 scaling to integrate the temperature-sensitive processes of exudation and degradation, enabling robust estimates of steady-state CPS under varying thermal conditions. Statistical results were visualized using ggplot2.

## Results

The degradation of 5-α-cyprinol sulfate (CPS) was investigated in a four-day experiment using autoclaved (Control) and non-autoclaved (NA) water treatments from two lakes: lake Fühlinger See (FS) and lake Reeser Meer Nord Erweiterung (RMNE). In the Control treatments (Fig. 1A), CPS concentrations showed no significant changes over time in either lake (FS: k=−0.0002, p=0.9286; RMNE: k=0.0025, p=0.4826, SI Table 1.). In the NA treatment (Fig. 1B), exponential models were fitted to assess CPS degradation rates. For Lake FS, the degradation constant (k) was 0.043 h⁻¹ (p< 0.01, indicating significant CPS degradation, SI Table 2.), corresponding to a 4.21% decay per hour (daily decay rate: 64.4 %, half-life: 16.1 h, Fig. 2). In contrast, Lake RMNE exhibited a higher degradation (constant of 0.081 h⁻¹ (p=0.0021, indicating significant CPS degradation, SI Table 2), corresponding to a 7.78% decay per hour (daily decay rate: 85.7 %, half-life: 8.6 h) (Fig. 2).

**Fig. 1:**
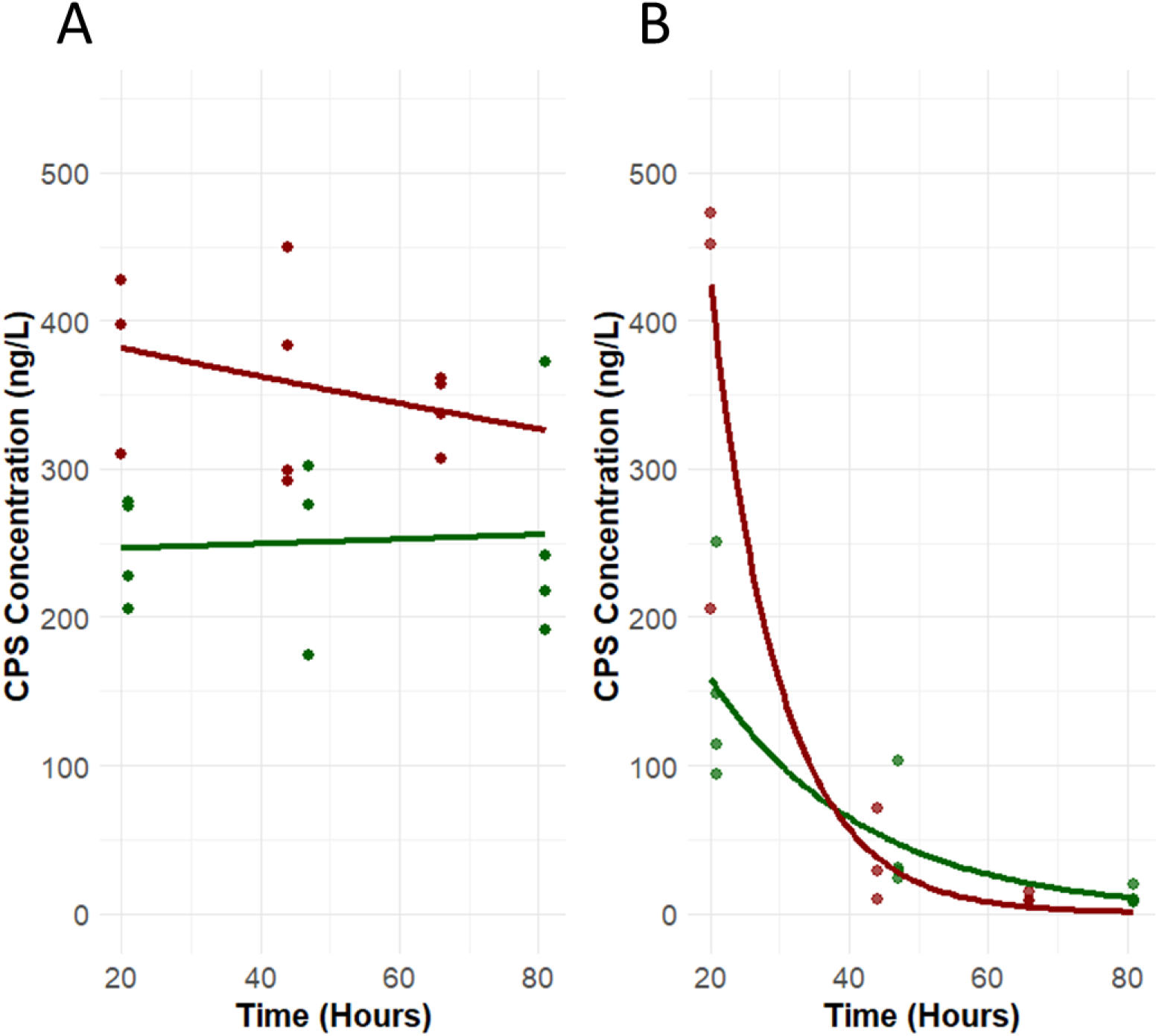
5α-cyprinol sulfate (CPS) concentration (ng/L) over time (hours) in lake water that was incubated indoors with and without initial autoclaving. Water samples originated from two lakes, FS (dark green) and RMNE (dark red), sampled in June 2024. (A) CPS concentration in autoclaved lake water (Control) with observed data points and fitted exponential decay models overlaid, showing no degradation over time in either FS (k = −0.0002, p = 0.9286) or RMNE (k = 0.0025, p = 0.4826) control treatments. (B) CPS concentrations in non-autoclaved lake water (NA) with observed data points and fitted exponential decay of CPS with modeled degradation curves overlaid on individual data points. Degradation rates (k) were calculated from the modeled degradation curves for RMNE (k=0.081, p=0.0021*; N*(*t*)=1651.21·*e*^-0.081·t^, *N* = CPS concentration at timepoint *t* in hours) and FS (k=0.043, p<0.01*; N*(*t*)=325.82·*e*^-0.043·t^, *N* = CPS concentration at timepoint *t* in hours). Each data point represents an individual measurement, and curves represent the modeled fit. N = 4 replicates for all treatments.

**Fig. 2:**
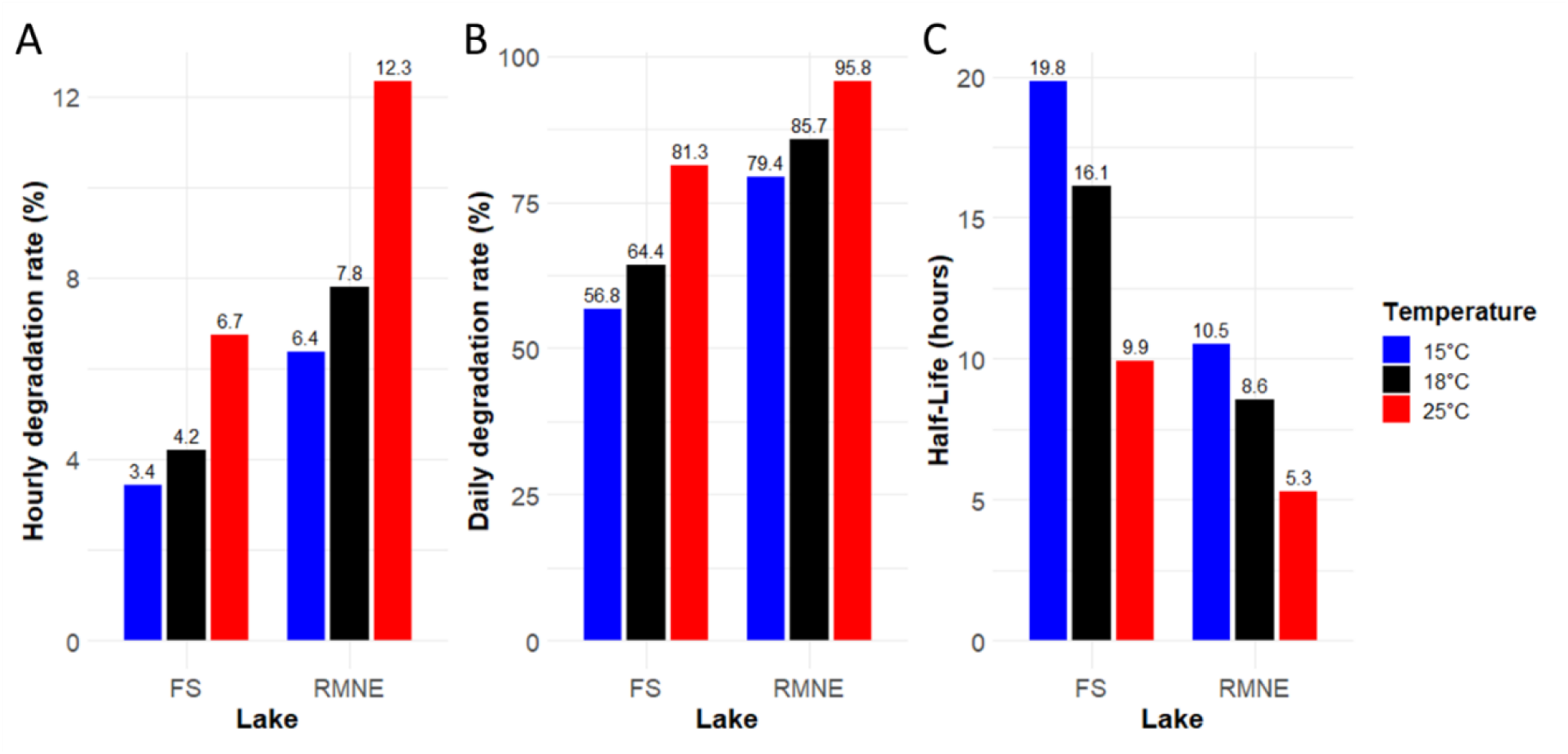
Hourly degradation rates (%), daily degradation rates (%), and half-life (hours) of 5α-cyprinol sulfate (CPS) at different temperatures in two lakes (FS and RMNE) in non-autoclaved lake water. **(A)** Percentage of CPS degraded per hour. **(B)** Percentage of CPS degraded per day. **(C)** CPS half-life in hours. Bars represent modeled values at 15°C (blue) and 25°C (red), calculated using Q_10_ scaling based on bacterial cell multiplication (BCM) as described by Simon and Wünsch (1998), alongside determined values at 18°C (black) derived from experimental data (n = 4). Percentage values are displayed above the bars for clarity.

The modeled impact of temperature and microbial activity on CPS degradation is shown in Figures 2 and SI Figure 1. CPS degradation in both lakes followed clear temperature-dependent trends, with higher temperatures leading to higher degradation rates, as predicted by Q10 scaling based on bacterial cell multiplication (BCM, Q10 ∼2) as described by Simon and Wünsch (1998). In FS (Fig. 2, SI Fig. 1B), the observed degradation rate at 18°C falls between the modeled predictions for 15°C and 25°C. The predicted hourly degradation rate increased from 3.4% at 15°C to 6.7% at 25°C (Fig. 2A), corresponding to daily degradation rates of 56.8% and 81.3%, respectively (Fig. 2B). The half-life of CPS decreased notably from 19.8 hours at 15°C to 9.9 hours at 25°C (Fig. 2C). In RNME (Fig. 2, SI Fig. 1A), CPS degradation was faster overall, with hourly degradation rates of 6.4% at 15°C, increasing to 12.3% at 25°C (Fig. 2A). This translated to daily degradation rates of 79.4% and 95.8% (Fig. 2B), with half-life values declining from 10.5 hours at 15°C to 5.3 hours at 25°C (Fig. 2C).

The CPS concentrations at timepoint 0 h were not directly measured but were estimated using the model developed for data from 20–80 h. This model, fitted to the observed CPS degradation rates, was used to calculate the initial concentrations by extrapolating back to 0 h. In FS, the predicted CPS concentration at 0 h was 325.82 ng/L (SI Fig. 1B), while in RNME, it was significantly higher at 1651.21 ng/L (p=0.0069) (SI Fig. 1A). These estimates provide valuable insight into the initial in-situ CPS concentrations and their differences between the two lakes.

In order to determine rates of CPS exudation by fish, a controlled laboratory experiment was performed with different amounts of food being fed to individual fish (*Rutilus rutilus*) kept in aquaria. CPS exudation (µg/fish) was measured over four consecutive days, and the cumulative amount of exuded CPS was plotted against the food amount (g) (Fig. 3). CPS-exudation rates deviated from normality (Shapiro–Wilk W = 0.40, p < 0.001), yet variances did not violate the assumption of homogeneity across food treatments (Levene F₍_25_, _80_₎ = 1.28, p = 0.20). A multiple linear regression model confirmed food amount as a significant predictor of CPS exudation (p<0.001), while fish size (i.e. body mass) had no significant effect (p=0.268). As fish size had no significant effect, it was excluded from the linear model, and only food amount was included as a predictor of CPS exudation. This led to a simplified model based solely on food amount. While the total CPS exudation per fish over 96 hours (Fig. 3) accounts for the full digestion process across different fish sizes, we calculated exudation rates per hour to better capture the direct relationship between food intake and CPS release dynamics. Since a rate rather than a total value was required for further integration into the model, the CPS exudation rate per hour was derived by fitting a simplified model: *y=−0. 8164+64.4952x*, where x represents the food amount (g) and y is the predicted CPS exudation (ng/h per fish). This model shows that for every additional gram of food, CPS exudation increased by approximately 64.5 ng/h per fish (p<0.001).

**Fig. 3:**
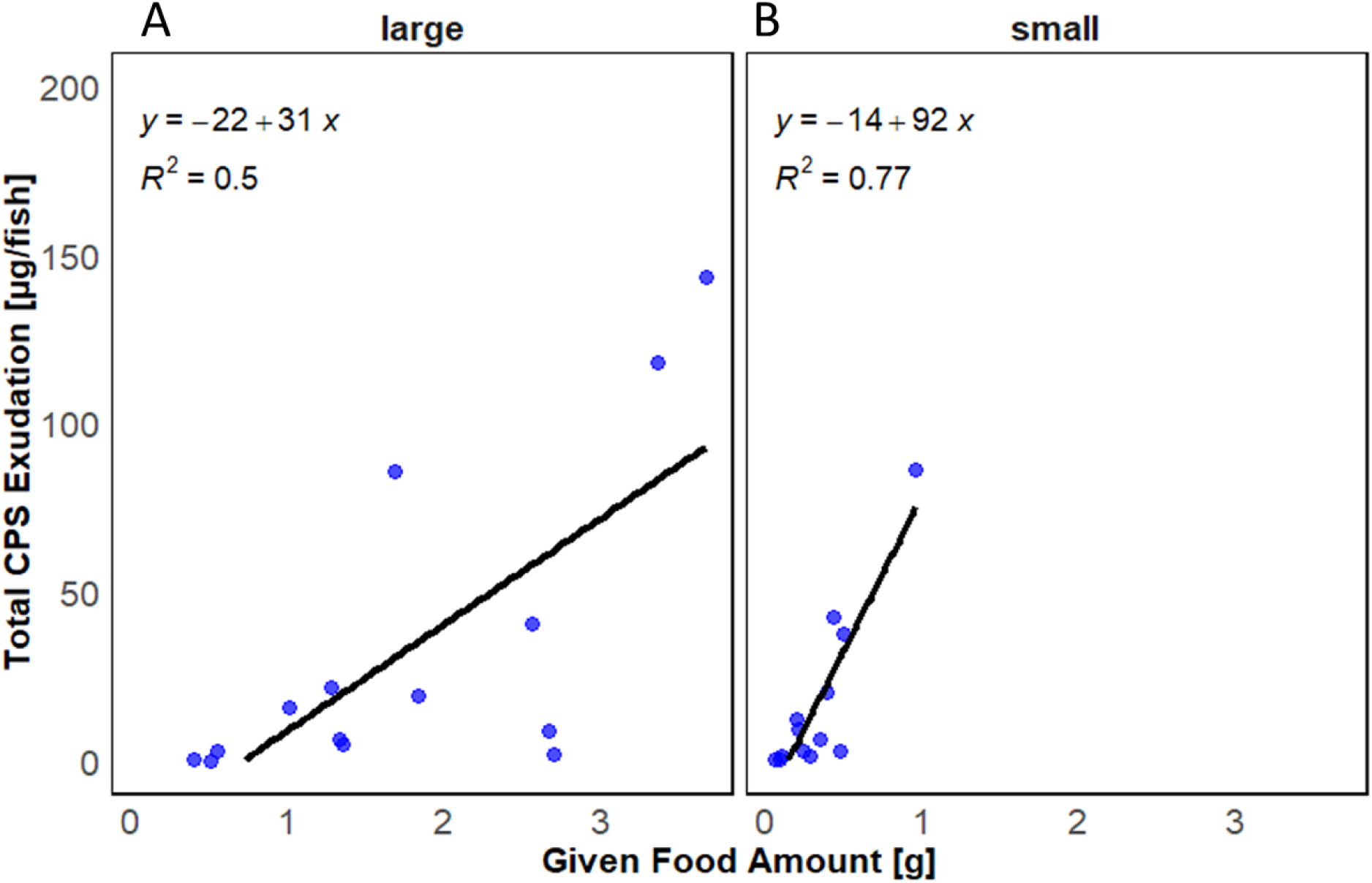
Relationship between 5α-cyprinol sulfate (CPS) exudation (µg/fish) and food amount (g) for two fish size categories. **(A)** Large fish (mean body mass 89.9 ± 13.5 g). **(B)** Small fish (mean body mass 18.2 ± 6.7 g). The x-axis represents the given food amount (g), and the y-axis indicates the total CPS exudation (µg/fish) 96 hours after the feeding event. Linear regression lines are fitted to the data, with corresponding equations and R² values provided in each panel to illustrate the fit of the regression models.

The seasonal dynamics of CPS concentrations and *Daphnia* abundances in RMNE were analyzed from March to September 2023. CPS concentrations (Fig. 4A) were below detection limit in March and September and ranged from 40 ng/L up to a peak concentration of 900 ng/L in August. Additionally, the seasonal temperature profile (Fig. 4B) revealed a progressive increase from March to August, peaking at 25°C, while water transparency measured as Secchi depth showed an inverse trend with reduced transparency during the warmer months. *Daphnia* ssp. were not detectable in March and nearly absent in April (0.3 Ind/L) (Fig. 5). Relative abundances increased sharply in May (37.1 Ind/L) peaked in June (80 Ind/L), and declined sharply in July (5 Ind/L) and August (2.3 Ind/L), with a resurgence in September (54.2 Ind/L).

**Fig. 4:**
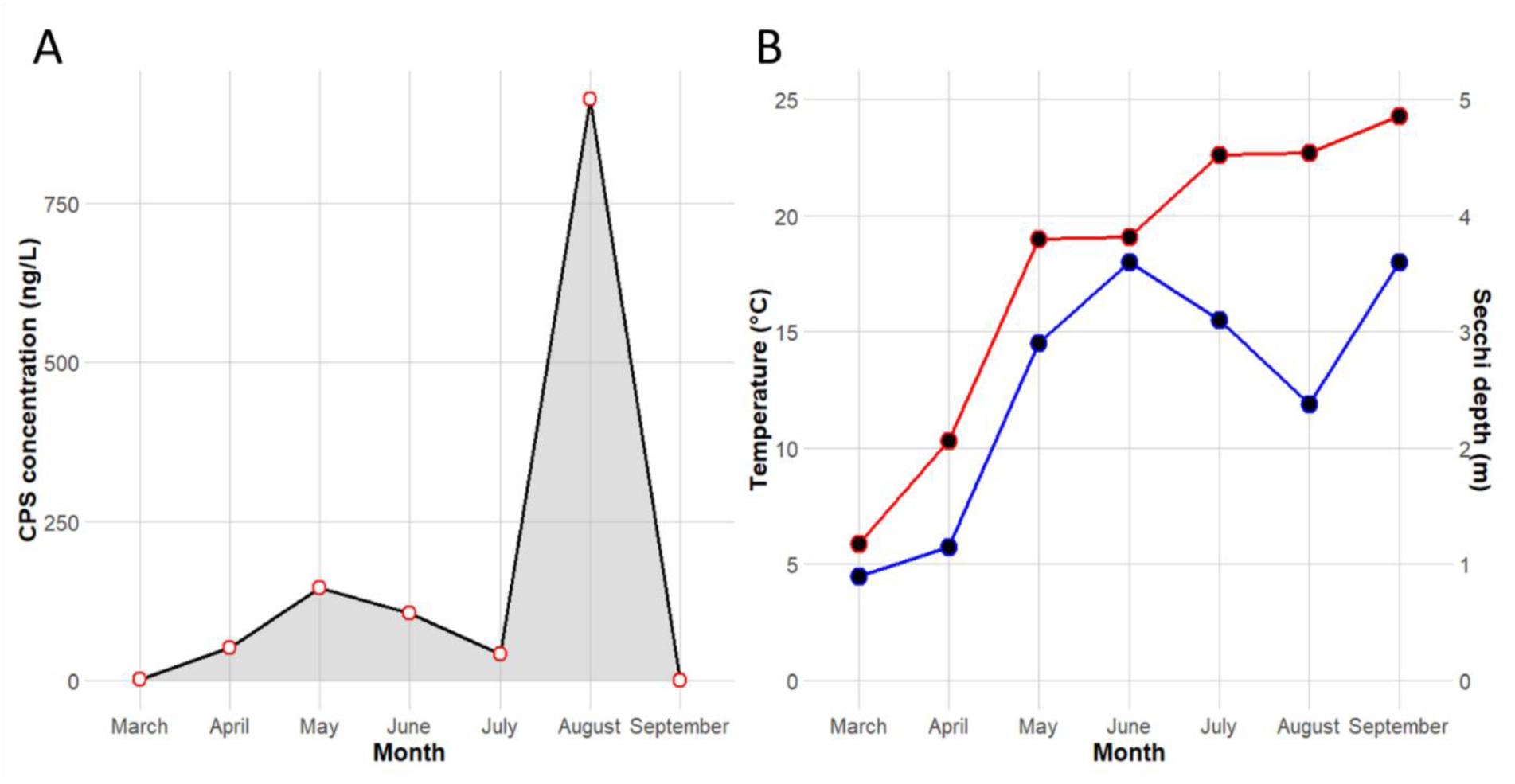
Seasonal dynamics of 5α-cyprinol sulfate (CPS) concentrations, water temperature, and Secchi depth in lake Reeser Meer Nord Erweiterung (RMNE) during 2023. **(A)** CPS concentrations (ng/L) from March to September, with the shaded area highlighting the seasonal trend. Points represent mean CPS concentrations derived from technical replicates (n = 4). **(B)** Temperature profile of the epilimnion (°C, red line) and Secchi depth (m, blue line) over the same period, illustrating seasonal variations in water transparency.

**Fig. 5:**
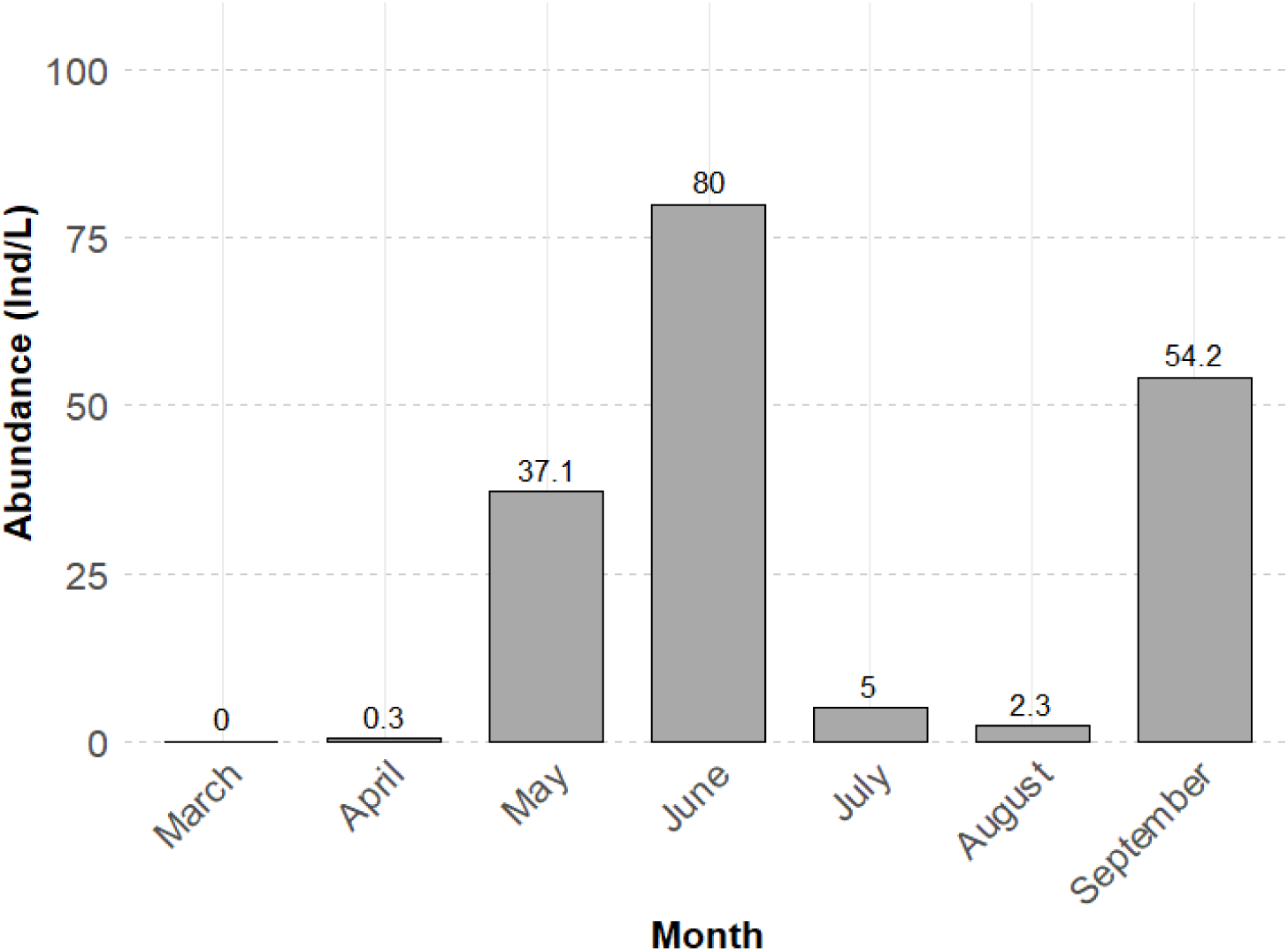
Abundance (Ind/L) of *Daphnia* in lake Reeser Meer Nord Erweiterung (RMNE) from March to September 2023. The values represent combined data from both the hypolimnion and epilimnion. Abundance is expressed in individuals per liter (Ind/L) for each month, with values displayed directly above each bar. (Data source: J. Borcherding, K. Scharnweber; pers. comm.)

Modeled CPS exudation per fish in lake RMNE showed a gradual increase from March to July, peaking in August, followed by a decline in September (Fig. 6). The model for CPS exudation incorporates not only the seasonal temperature profile but also daily zooplankton food rations of fish based on the study by Persson (1983). Including daily food rations in the model allows us to capture the combined effects of seasonal variations in food availability and metabolic processes, providing a more comprehensive understanding of CPS dynamics. By integrating both exudation and degradation processes, we aimed to investigate the balance between CPS release and microbial breakdown, enabling a more accurate prediction of in-situ CPS concentrations under shifting environmental conditions.

**Fig. 6:**
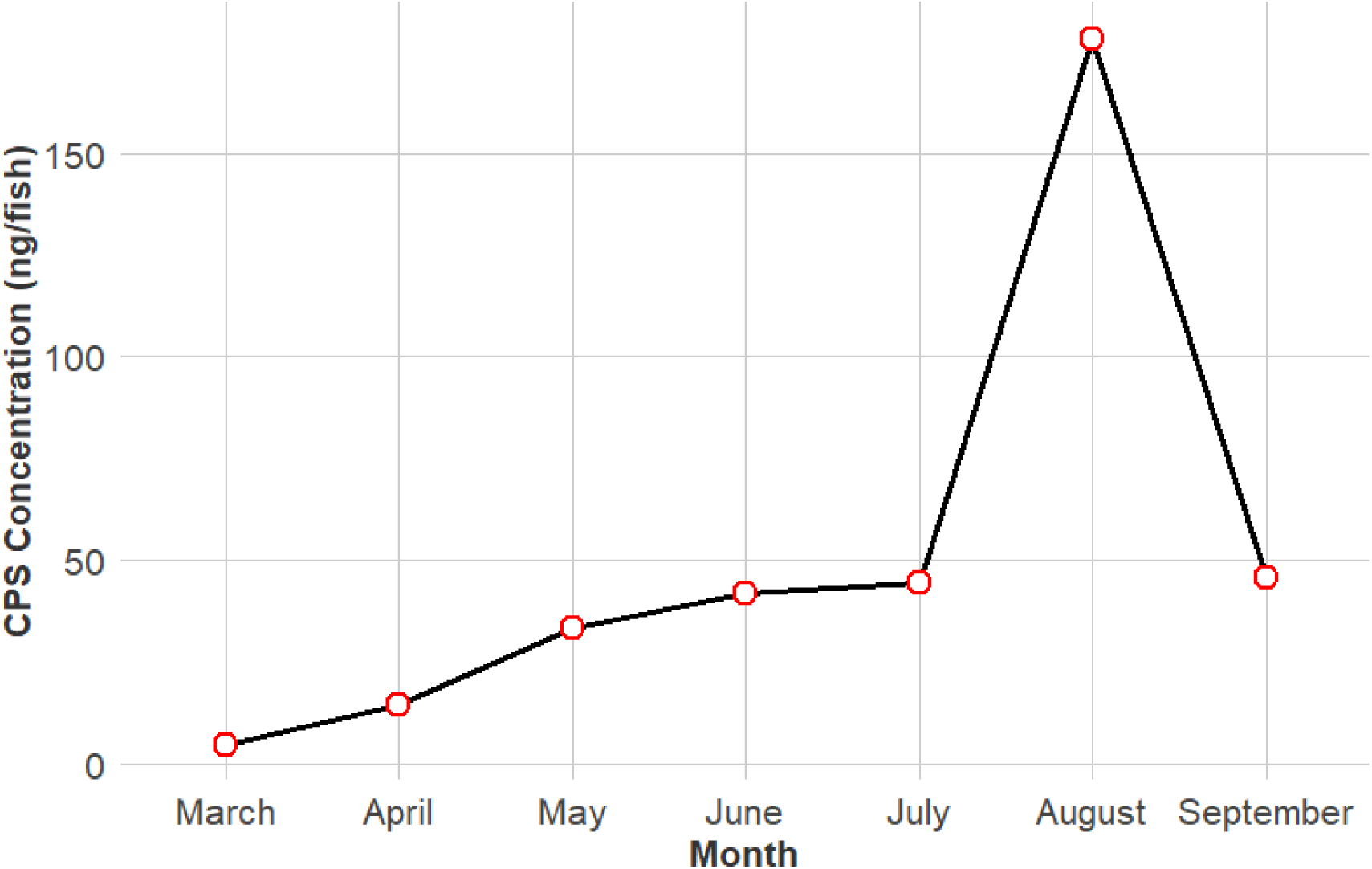
Modeled net exudation of 5α-cyprinol sulfate (CPS) per individual fish (ng/fish) from March to September. The model incorporates monthly temperature data from lake Reeser Meer Nord Erweiterung (RMNE) in 2023 and food consumption data derived from Persson (1983). The model accounts for temperature-dependent exudation and degradation rates alongside food intake, illustrating seasonal metabolic dynamics influenced by environmental conditions.

## Discussion

### Microbial CPS degradation

The bacterial degradation of the diel vertical migration (DVM)-inducing kairomone exuded by cyprinids was first demonstrated in the laboratory study by Loose et al. (1993). However, this proof-of-concept study did not provide insights into the relevance of this process under natural conditions. More recent mesocosm experiments (Ahlers & Hahn et al. 2024) indicated potential in-situ CPS degradation, though microbial involvement was not conclusively established.

In our study, we provide robust evidence that microbial degradation is the primary driver of CPS breakdown in both lakes that we have investigated here, as demonstrated by the absence of degradation in autoclaved control samples. The determined half-lives at 18°C were 8.6 hours in RMNE and 16.1 hours in FS, indicating that, on average, approximately 85.7% and 64.4% of the CPS compound is degraded daily, respectively. These differences in degradation rates suggest variations in microbial community composition, activity, or environmental conditions influencing CPS dynamics. The faster degradation in RMNE likely reflects a more specialized microbial community, potentially adapted to utilizing CPS as a carbon source while mitigating its inherent toxicity due to its amphiphilic nature (Philipp 2011; Feller et al. 2021; Müller et al. 2021). Furthermore, higher initial CPS concentrations in RMNE may have stimulated microbial activity through substrate-induced enzymatic activation, a well-documented phenomenon in microbial ecology (Boetius and Lochte 1994; Hernández et al. 2012; Allison et al. 2014). These results underscore the intricate relationship between microbial communities and environmental factors in regulating CPS degradation.

### Temperature as a key modulator of microbial degradation

Temperature has emerged as a critical factor influencing degradation rates, as demonstrated by the Q10 modeling approach. Bacteria play a key role in dissolved organic carbon degradation, with studies showing that microbial activity drives a significant amount of carbon breakdown in freshwater systems (Tranvik et al. 2009; Kawasaki et al. 2013). For this reason, the Q10 values used to model CPS degradation rates were derived from bacterial growth rates under varying temperature conditions, as reported by Simon and Wünsch (1998) and Beveridge et al. (2010). Temperature affects microbial metabolism and community dynamics by regulating cellular processes such as enzyme activity, metabolic rates, and trophic interactions (Beveridge et al. 2010; Moon et al. 2023). These effects, in turn, shape broader ecosystem functions, emphasizing the critical role of temperature in microbial-mediated degradation processes. For instance, in RMNE, degradation rates increased from 6.4% at 15°C to 12.3% at 25°C per hour, with the half-life of the CPS-molecule effectively halving with this increase of temperature. This rapid degradation of CPS mirrors findings by Mendelski et al. (2019), who observed complete degradation of the bile salt chenodeoxycholic acid in soil environments within 20 hours at room temperature. The rapid microbial breakdown observed in our study challenges previous assumptions about persistence of bile salts (and thus as well of CPS) in aquatic environments, emphasizing the transient nature of this infochemical. (Hofmann and Mysels 1987; Buchinger et al. 2014; Wyatt 2014).

### Linking CPS exudation to feeding activity

Zooplankton populations fluctuate seasonally, strongly influencing the feeding rates of zooplanktivorous fish (Persson 1983). Persson’s study demonstrated significant seasonal fluctuations in fish food intake, with higher consumption rates observed during warmer months due to increased metabolic demands and greater resource availability. Our exudation experiments demonstrated that CPS release is directly proportional to food intake. According to the Plankton Ecology Group (PEG) model (Sommer et al. 1986; Sommer et al. 2012) fish feeding rates closely follow seasonal shifts in prey availability, which, according to our results, in turn will influence CPS excretion. This suggests that natural CPS release is governed primarily by feeding intensity rather than by fish size or body mass. This challenges traditional metabolic scaling principles, which propose that excretion rates scale non-linearly with body size (Vanni and Gephart 2011; Morgan and Hicks 2013).

Instead, our findings indicate that CPS excretion follows a different pattern, likely driven by feeding frequency rather than metabolic constraints. Bile compounds are secreted into the gut lumen in response to food intake through complex physiological mechanisms (Maldonado-Valderrama et al. 2011; Buchinger et al. 2014; Di Ciaula et al. 2017). CPS constitutes the major bile compound in cyprinid fish (Hagey et al. 2010), and consequently, periods of increased feeding, such as during zooplankton blooms, are expected to result in significant CPS excretion peaks. The constancy of factorial aerobic scope (FAS) across fish sizes suggests that metabolic efficiency remains tightly regulated regardless of body size, further supporting the conclusion that food intake is the primary driver of CPS release (Huang et al. 2013).

Our results further demonstrate that, while larger fish can ingest greater amounts of food and thus excrete higher absolute quantities of CPS, size-normalized excretion rates are remarkably similar across different fish sizes. This finding aligns with Persson (1983), who observed that older roach have higher predicted daily food rations than younger ones, reinforcing the conclusion that food consumption is the primary driver of CPS exudation. As a consequence, seasonal dynamics of planktonic organisms play a fundamental role in shaping CPS exudation patterns in lakes.

CPS has been demonstrated to induce two types of defenses in *Daphnia*, i.e. DVM and morphological changes (Hahn et al. 2019; Hahn & von Elert 2022; Ahlers & Hahn et al. 2024). We here show that the release of CPS by cyprinid fish is linearly related to the ingested food amount, which is in accordance with the function of bile salts as emulsifiers of lipids in the gut lumen. Accordingly, we expect that the identity of the prey is less important, so that the rate of kairomone release would not change greatly if a fish were fed *Daphnia* instead of mosquito larvae. However, it should be noted that for inducible changes in life history of *Daphnia*, for which the chemical nature of the kairomone released by fish is yet unknown (Von Elert & Stibor 2006), it has recently been demonstrated that the release of the kairomone depends on that the prey is phylogenetically closely related to *Daphnia* (Gu et al 2023).

### Seasonal interplay of CPS and environmental factors

The seasonal dynamics of CPS concentrations in RMNE provide valuable insights into the interconnected roles of fish feeding, environmental conditions, and prey availability. CPS concentrations fluctuated throughout the season (from 40 ng/L up to a peak concentration of 900 ng/L), with a pronounced peak in August, likely driven by increased fish feeding activity during this period (Sommer et al. 2012).

Furthermore, experimental studies have demonstrated that CPS concentrations must exceed specific thresholds to induce behavioral and morphological responses in *Daphnia* species. Specifically, concentrations above 4-17 µg/L were required to trigger diel vertical migration (DVM) in *Daphnia longispina* in a mesoscosm experiment (Ahlers and Hahn et al. 2024) and in *D. magna* in an indoor setup DVM was induced already with 53 ng/L (Hahn et al. 2019). Morphological defenses such as helmet and spine elongation in the invasive species *Daphnia lumholtzi* have been observed at even lower concentrations, with significant changes occurring already at 5.31 ng/L (Hahn & von Elert, 2022). These findings highlight the ecological relevance of CPS exudation, demonstrating that even minimal concentrations can elicit profound biological effects.

When comparing seasonal CPS profiles with *Daphnia* abundance data, it becomes evident that the latter alone does not fully explain the observed CPS dynamics in RMNE. Although *Daphnia* populations peaked in June, CPS concentrations remained relatively low, suggesting that fish predation on *Daphnia* alone cannot fully account for kairomone levels. As *Daphnia* abundances declined in July and August, fish likely shifted their diet to alternative prey, such as *Chaoborus*, other zooplankton, or phytoplankton, as suggested by previous studies (Persson 1982; Persson 1983; Brabrand 1985; Haertel and Eckmann 2002). During this time of the year, young fish have increased in body size and gape size, enabling them to target larger prey items, further diversifying their diet. Therefore, it seems reasonable to assume that this dietary flexibility allows fish to sustain CPS excretion even when *Daphnia* populations are reduced.

Interestingly, in August, water transparency reached its seasonal minimum, coinciding with the peak in CPS concentrations. The simultaneous occurrence of high CPS concentrations and decreased water transparency supports the hypothesis that elevated fish activity, particularly in response to increased temperatures, intensifies predation pressure on zooplankton, indirectly leading to higher phytoplankton biomass and reduced water clarity.

### Rapid turnover of CPS in aquatic systems

One of the key findings of this study is the exceptionally rapid turnover of CPS in aquatic systems. Despite degradation rates of up to 85.7% per day in RMNE, in-situ CPS concentrations remained relatively stable, indicating a dynamic equilibrium between degradation and continuous exudation from cyprinid fish. At 18°C, over 85% of CPS molecules are degraded daily, yet the steady concentrations suggest a finely tuned balance capable of rapidly responding to environmental fluctuations.

This high turnover challenges the previously held assumption of CPS stability in aquatic environments and instead suggests that in-situ concentrations are highly dynamic and responsive to biotic and abiotic changes. Previously, CPS was considered a reliable indicator of predation risk, with DVM amplitudes in *Daphnia* increasing in response to CPS concentrations (Hahn and von Elert 2022). However, our findings suggest a more nuanced perspective: CPS concentrations do not solely reflect predator abundance but are also strongly influenced by predator feeding activity and microbial degradation. This underscores the need to reinterpret CPS as a highly dynamic chemical signal, modulated by ecological and environmental conditions rather than being a stable proxy for predation pressure alone.

### Advancing predictive models for CPS and fish dynamics

The model for prediction of net CPS exudation represents a first step towards integrating biotic and abiotic factors into a predictive framework. Notably, the model successfully reflects the seasonal pattern of the observed CPS concentrations in RMNE, underscoring the critical role of fish feeding activity in shaping the CPS dynamics over time. However, the pronounced decline in CPS concentrations observed in June cannot be accounted for by the current model, which strongly suggests that additional factors, such as prey availability, shifts in fish diet, or abiotic influences, must be incorporated to improve the model’s predictive accuracy. Furthermore, the current model does not fully account for the dynamic nature of CPS concentrations, as it primarily reflects long-term (monthly) shifts in biotic and abiotic factors. The model may overlook short-term (daily) fluctuations that could play a crucial role in CPS turnover and ecosystem signaling. Incorporating variables like diel feeding rhythms, prey-specific exudation rates, and broader ecosystem interactions would offer a more comprehensive understanding of CPS dynamics in aquatic systems. Additionally, the model could serve as a useful tool for indirectly estimating fish populations, addressing the scarcity of reliable data on fish abundance in natural environments. By integrating key factors such as temperature, food intake, and exudation patterns, this model has the potential to bridge critical knowledge gaps and provide a novel framework for understanding fish-mediated ecological processes in complex aquatic ecosystems.

## Acknowledgements

We extend our sincere gratitude to Jost Borcherding, Ann-Marie Waldvogel, Kristin Scharnweber and Alexander Popov from the Ökologische Forschungsstation Rees for their invaluable expertise, support, and provision of *Daphnia* abundance data and lake water samples throughout the season at Lake RMNE. We also thank Patrick Fink and Andrea Hoff from the UFZ Water Analytics Laboratory (GEWANA) for their assistance in conducting the phosphorus measurements. We thank Marie Olbrich who performed the indoor fish exudation experiments. We are especially grateful to Johannes for his expert statistical advice, which greatly improved our analyses. This research was generously supported by the German Science Foundation (DFG) through a grant awarded to EVE (EL 179/16-1). Open Access funding was partly provided by the University of Cologne and facilitated through Projekt DEAL.

---

**SI Fig. 1:**
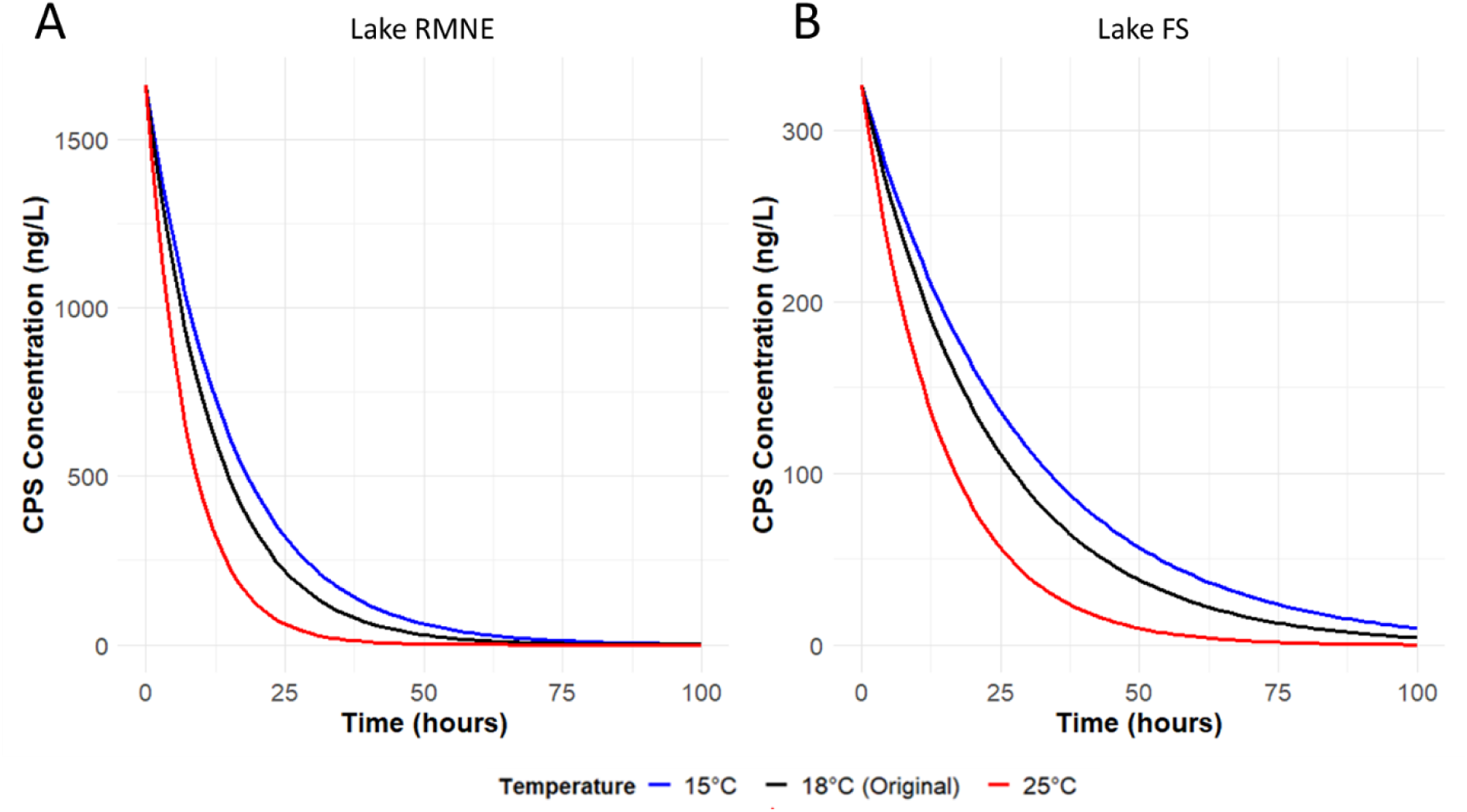
Modeled exponential decay of 5α-cyprinol sulfate (CPS) concentrations over time (ng/L) for non-autoclaved lake water from two lakes sampled in June 2024. (A) CPS concentrations in lake RMNE. (B) CPS concentrations in lake FS. The black line represents modeled CPS concentrations at 18°C based on determined data. The blue and red lines represent modeled CPS concentrations adjusted to 15°C and 25°C, respectively, using Q10 scaling based on bacterial cell multiplication (BCM) as described by Simon and Wünsch (1998). The modeled curves span the time interval from 0 to 100 hours.

**SI Table 1:** Results of a linear mixed-effects model (lme) used to analyze log-transformed CPS concentrations of autoclaved water (Control) over time in degradation experiments.

| *Effect* | *Estimate* | *Std. Error* | *t-value* | *p-value* | *Interpretation* |
| --- | --- | --- | --- | --- | --- |
| *(Intercept)* | 5.493 | 0.127 | 43.415 | <0.001 | Log-transformed initial CPS concentration for FS (reference lake). |
| *time* | 0.0002 | 0.002 | 0.091 | 0.929 | No significant degradation rate (k) for FS. |
| *RMNE* | 0.490 | 0.188 | 2.604 | 0.017 | RMNE has a significantly higher initial CPS concentration compared to FS. |
| *time:RMNE* | -0.003 | 0.004 | -0.716 | 0.483 | No significant difference in degradation rates between RMNE and FS. |

**SI Table 2:**
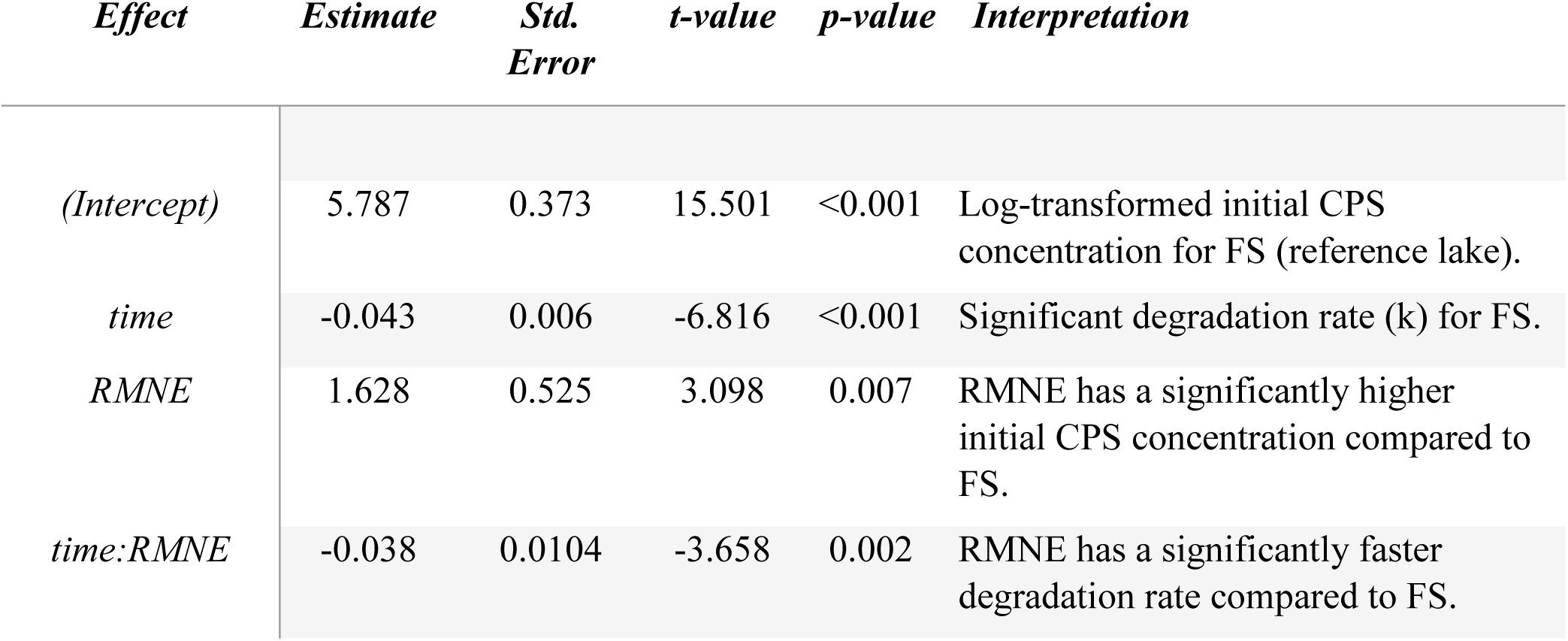
Results of a linear mixed-effects model (lme) used to analyze log-transformed CPS concentrations of non-autoclaved water (NA treatment) over time in degradation experiments.

